# Crowdsourcing biocuration: the Community Assessment of Community Annotation with Ontologies (CACAO)

**DOI:** 10.1101/2021.04.30.440339

**Authors:** Jolene Ramsey, Brenley McIntosh, Daniel Renfro, Suzanne A. Aleksander, Sandra LaBonte, Curtis Ross, Adrienne E. Zweifel, Nathan Liles, Shabnam Farrar, Jason J. Gill, Ivan Erill, Sarah Ades, Tanya Z. Berardini, Jennifer A. Bennett, Siobhan Brady, Robert Britton, Seth Carbon, Steven M. Caruso, Dave Clements, Ritu Dalia, Meredith Defelice, Erin L. Doyle, Iddo Friedberg, Susan M.R. Gurney, Lee Hughes, Allison Johnson, Jason M. Kowalski, Donghui Li, Ruth C. Lovering, Tamara L. Mans, Fiona McCarthy, Sean D. Moore, Rebecca Murphy, Timothy D. Paustian, Sarah Perdue, Celeste N. Peterson, Birgit M. Prüß, Margaret S. Saha, Robert R. Sheehy, John T. Tansey, Louise Temple, Alexander William Thorman, Saul Trevino, Amy Cheng Vollmer, Virginia Walbot, Joanne Willey, Deborah A. Siegele, James C. Hu

## Abstract

Experimental data about known gene functions curated from the primary literature have enormous value for research scientists in understanding biology. Using the Gene Ontology (GO), manual curation by experts has provided an important resource for studying gene function, especially within model organisms. Unprecedented expansion of the scientific literature and validation of the predicted proteins have increased both data value and the challenges of keeping pace. Capturing literature-based functional annotations is limited by the ability of biocurators to handle the massive and rapidly growing scientific literature. Within the community-oriented wiki framework for GO annotation called the Gene Ontology Normal Usage Tracking System (GONUTS), we describe an approach to expand biocuration through crowdsourcing with undergraduates. This multiplies the number of high-quality annotations in international databases, enriches our coverage of the literature on normal gene function, and pushes the field in new directions. From an intercollegiate competition judged by experienced biocurators, Community Assessment of Community Annotation with Ontologies (CACAO), we have contributed nearly 5000 literature-based annotations. Many of those annotations are to organisms not currently well-represented within GO. Over a ten-year history, our community contributors have spurred changes to the ontology not traditionally covered by professional biocurators. The CACAO principle of relying on community members to participate in and shape the future of biocuration in GO is a powerful and scalable model used to promote the scientific enterprise. It also provides undergraduate students with a unique and enriching introduction to critical reading of primary literature and acquisition of marketable skills.

**Significance Statement:** The primary scientific literature catalogs the results from publicly funded scientific research about gene function in human-readable format. Information captured from those studies in a widely adopted, machine-readable standard format comes in the form of Gene Ontology annotations about gene functions from all domains of life. Manual annotations based on inferences directly from the scientific literature, including the evidence used to make such inferences, represents the best return on investment by improving data accessibility across the biological sciences. To supplement professional curation, our CACAO project enabled annotation of the scientific literature by community annotators, in this case undergraduates, which resulted in contribution of thousands of validated entries to public resources. These annotations are now being used by scientists worldwide.

## Introduction

Biocuration captures information from the primary literature in a computationally accessible fashion. The biocuration process generates annotations connecting experimental data with unique identifiers representing precisely defined ontology terms and logical relationships. While the majority of existing annotations are computational predictions built on knowledge from human biocuration, manually curated annotations from published experimental data are still the gold standard for functional annotations(1). Universal access to well-curated databases, such as UniProt and those maintained by model organism consortia, allows scientists worldwide to leverage computational approaches to solve pressing biological problems. New insights on complex cellular processes such as autophagy, cell polarity, and division can be clarified after assessing relationships in curated data(2–4). The Gene Ontology (GO, http://geneontology.org/) is an evolving biocuration resource that provides the framework for capturing attributes of gene products within three aspects or main branches: biological process, molecular function, and cellular component(5, 6). Importantly, connections can be made between model organism genes and human genes with comprehensive GO coverage(7). Additionally, using GO data generates testable hypotheses in areas with little direct experimentation(8–10). Application to high-throughput and systems biology, for instance, has led to insights and better methods for identification and analysis of the genes involved in cardiac and Alzheimer’s disease(11, 12).

Without question GO is a critical scientific resource, but manual annotation is an extremely labor-intensive process(13, 14). The pace at which the information is generated in the literature exceeds the capacity of professional biocurators to perform manual curation and the willingness of funding agencies to pay for a larger biocurator labor force(15). Although the general Swiss-Prot protein database (https://www.uniprot.org/) model is one example that has kept up with manual and automated annotations, many fields are limited by low numbers of trained personnel and minimal participation, even by trained scientists(16, 17). The problem is most severe for communities studying organisms without a funded model organism database. Nevertheless, curation of the experimental literature from as many species as possible strengthens inference of function when there is substantial evolutionary conservation(18, 19). Several groups are developing tools to facilitate community engagement, such as the Gene Ontology Normal Usage Tracking System (GONUTS) site described here. These efforts stem from the realization that, while most scientists acknowledge the importance of data curation, it is hard to motivate individuals to volunteer their knowledge(20, 21). Spectacular successes include crowdsourcing the analysis of the 2011 Shiga-toxin producing *E. coli*(22), the solution of the structure of an HIV protease by the FoldIt player community(23), and science content within Wikipedia(24–27). In other cases, high-profile community annotation efforts have been less successful(28). Previously, Dr. James Hu was quoted in *Nature* describing the fundamental challenge for community participation in biocuration in terms of the traditional incentives for funding and promotion in academia(29).

Here, we describe the successful implementation over nearly a decade of a university instruction-based model resulting in nearly 5000 high-quality community annotations added to the GO database. This effort was motivated by the clear parallels between the foundational skills used in the professional biocuration field and the well-defined goals for undergraduate training(30). A professional GO biocurator creates gene annotations by finding relevant primary literature, extracting information about normal gene function from it, and entering that information using the controlled GO vocabulary into online databases(31). We hypothesized that university students, guided by their instructors, could accomplish similar tasks and perform community GO annotation while developing strong critical reading skills in a templated annotation task requiring rigorous reading of primary scientific literature. The GONUTS wiki platform (https://gowiki.tamu.edu/) was originally built as a framework for experts not familiar with GO to annotate from literature in their field. We leverage GONUTS to allow student structured GO annotation entry (Fig. 1)(32).

**Figure 1:**
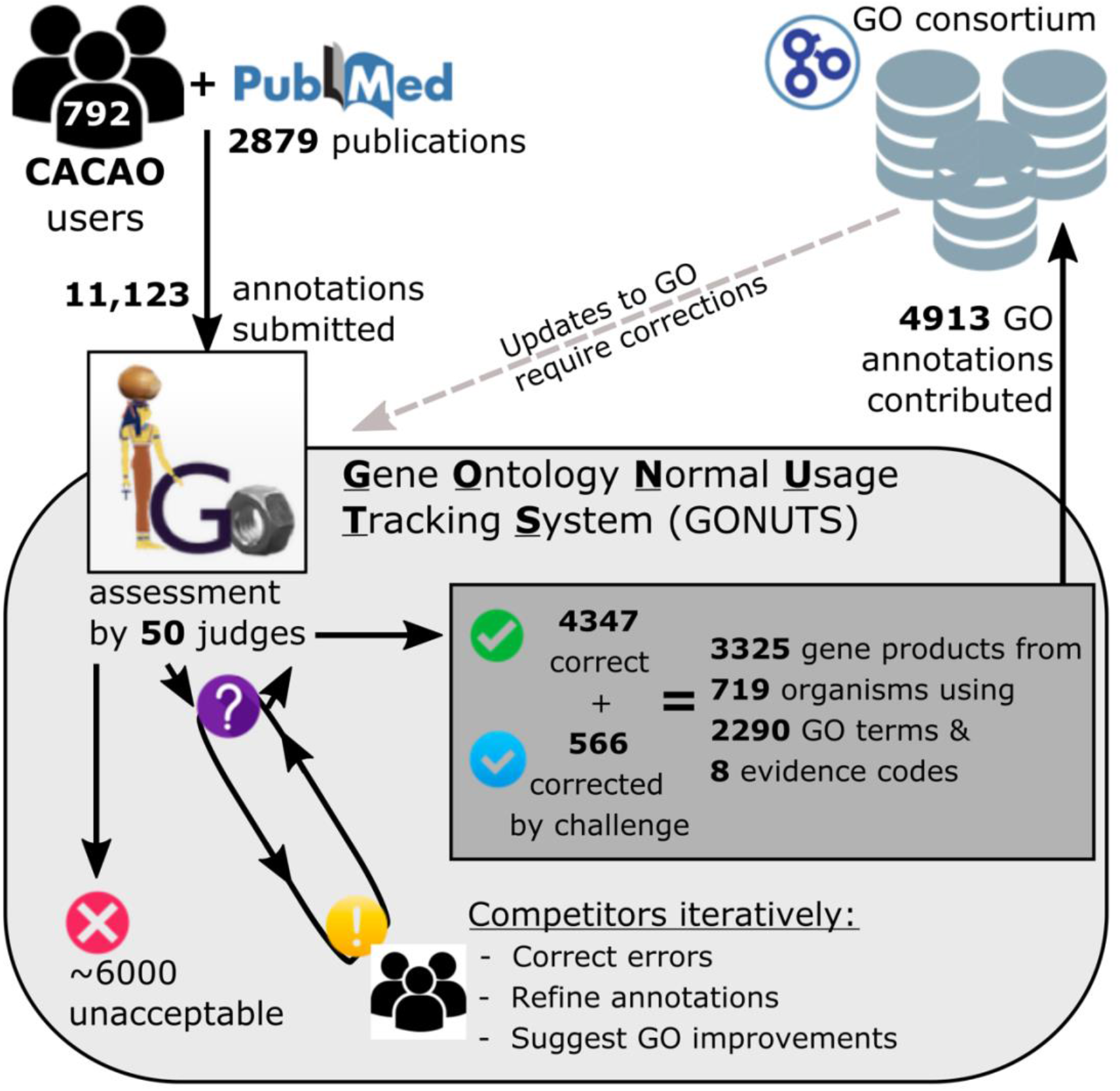
CACAO competitors contribute a large number of GO annotations. Overall CACAO contributions are summarized in the context of the workflow for quality control and submission to the GO Consortium. CACAO users consume the primary literature, collect information about normal gene functions from the paper study subjects, and capture the evidence and conclusions using the Gene Ontology. Those annotations are reviewed by trained judges and marked as unacceptable (red X), requiring changes (yellow !, or purple ? flagged for further review), or acceptable (green check, or blue check after correction) within the GONUTS framework. Competitors challenge entries and engage in peer review until an annotation is corrected or marked unacceptable. Fully vetted annotations are deposited into the public GO database maintained by professional biocurators and used by scientists worldwide. As required, CACAO-submitted annotations will be updated to reflect rearrangements and changes in GO.

## Results

### Sustainable community member contribution via an online intercollegiate competition

To address our need for broader participation and expansion beyond model organism databases, we initiated an intercollegiate competition based at Texas A&M University mainly for undergraduate students, called Community Assessment of Community Annotation with Ontologies (CACAO). The specifics of teaching practice are beyond the scope of this report (33). Here, we limit the discussion to details of the competition that are relevant to annotation. Teams of students (competitors) participate in the CACAO competition. Instructors (also called judges) assess all annotations entered by competitors for accuracy and completeness, then give feedback. Peer review by the competitors is incentivized by awarding points for challenges that correct an entry. Teams earn points only for correct annotations and challenges. The team with the highest points accumulated over the competition period wins. Vetted, high-quality annotations are then submitted to the GO Consortium database. CACAO quickly expanded, hosting 39 competitions over eight years including 23 colleges and universities, with 792 community annotators and 50 judges. After reading 2879 peer-reviewed journal articles, community members submitted 11,123 annotations to GONUTS (Fig. 1). Many were rejected, usually as unsuitable for GO annotation. Following careful review of each facet for every annotation submitted through online CACAO competitions, 4913 diverse annotations were added to the GO Consortium database after 2018 (Fig. 1). Those annotations are maintained as mandated by updates or changes in the ontology.

### Annotations generated through CACAO are diverse and novel

The 4913 annotations contributed through GONUTS have spanned all domains of life plus viruses, with the majority being skewed towards eukaryotes, in particular model organisms among the chordates (human, mouse, rat, *etc*.), *Streptophyta* (plants including *Arabidopsis*), and *Ascomycota* (such as budding yeast) (Fig. 2A). As only unique annotations are accepted, this demonstrates that community members can help fill the gaps left by professional biocurators working for model organism databases. Archaea and archaeal viruses are sparsely annotated in GO and represent the smallest fractions within our set, with only 24 and six annotations each, respectively. In contrast, 285 eukaryotic viruses are represented, and 45% of the viral annotations cover 384 viruses that infect bacteria (phages include *Siphoviridae*, *Myoviridae*, *Podoviridae*, and *Tectiviridae*). Nearly half of the 1000 annotations listed for bacterial viruses (phages) in QuickGO list CACAO as the source. Annotations for bacterial proteins make up only 5% of total GO annotations, but 30% of CACAO annotations. At the Order level, the top five bacterial categories (*Enterobacterales*, *Bacillales*, *Lactobacillales*, *Pseudomonales*, *Vibrionales*) are heavily studied Gram-negative and Gram-positive organisms of importance to microbiology research and the medical community. The microbial (virus and bacteria) entities herein described represent high genetic diversity and often serve as the basis for significant automated propagation to eukaryotic gene products. Thus, we conclude that not only do CACAO annotators fill gaps for model organisms, but also expand coverage to a wide array of otherwise poorly curated species.

**Figure 2:**
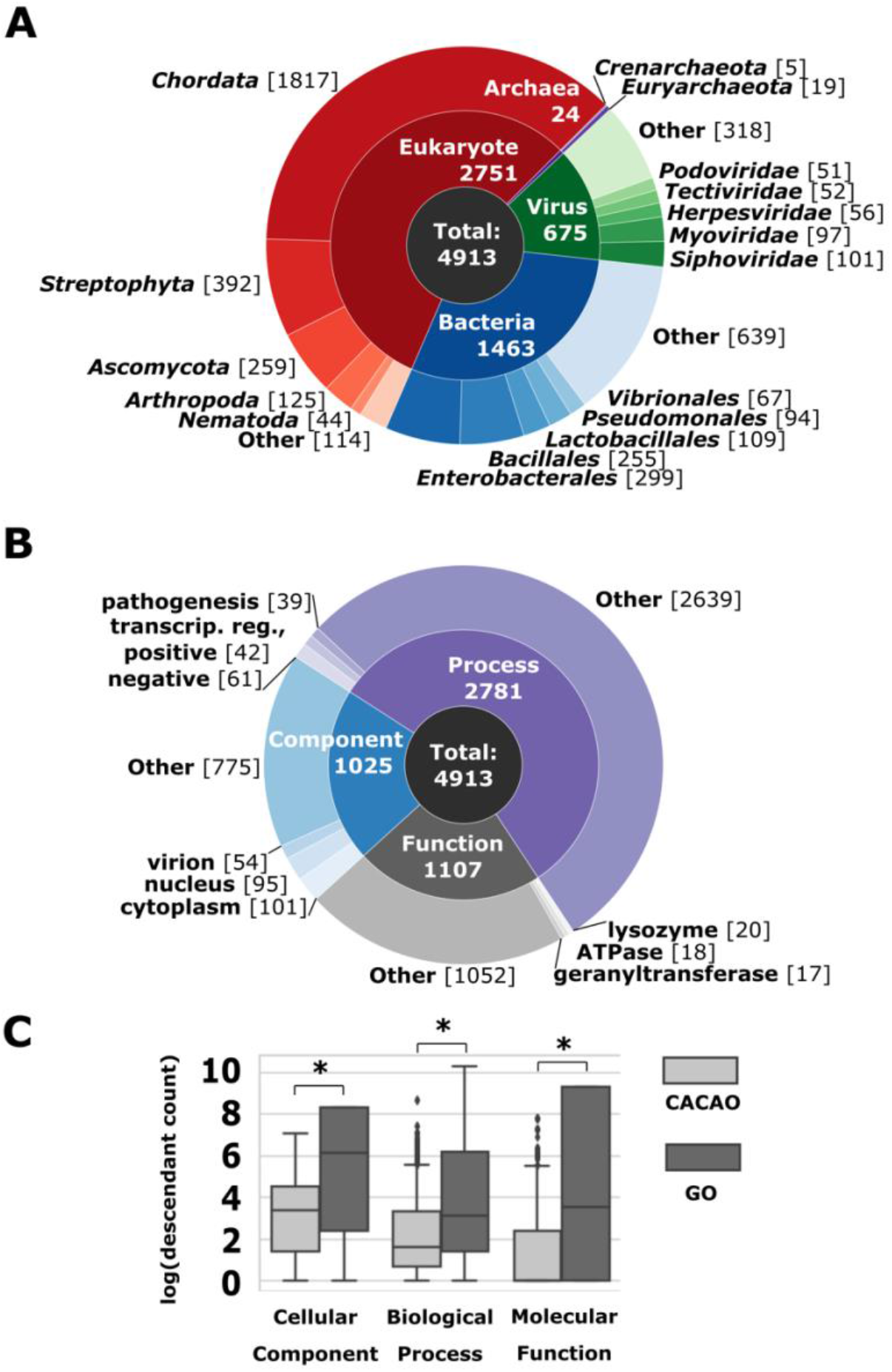
The GO annotations contributed by CACAO users are diverse and specific. A) Proteins annotated by CACAO users are depicted by species domain. The organisms most highly represented in each domain are displayed on the outer ring of the chart divided by the following rank: Phylum for eukaryotes and archaea, Order for bacteria, and Family for viruses. The number of GO annotations in each category is indicated in brackets. B) The distribution of GO terms used for CACAO annotations are graphed by aspect within the ontology. The top three terms within each aspect are labeled on the outer ring. For clarity, “activity” was dropped from each function term, and the process terms were abbreviated from “positive/negative regulation of transcription, DNA-templated” to “transcript. reg., positive or negative”. The number of GO annotations for each term is indicated in brackets. C) The descendant counts, corresponding to depth within the ontology, for CACAO annotations (n = 4913) and all other manual GO annotations through 2019 (n = 255,958) are graphed. Significant differences measured by the Mann-Whitney test with p<0.001 are marked with an *.

Interestingly, GO process terms are used in more than half of the CACAO annotations (Fig. 2B). The top three terms used within each aspect (Process, Component, and Function) are only a small proportion of the total for that branch, an indicator that community members annotate to a wide variety of terms. The cellular component terms are relatively general (nucleus), likely reflecting the ambiguity of experimental methods typically reported in papers. In contrast, the top process and function terms are near leaf-level, having few to no child terms, indicating specific annotations. To better understand the level of detail captured in annotations made by CACAO users, we used GOATOOLS, a python package developed by Klopfenstein *et al.* for representing where terms fall within the ontology hierarchical graph(34). Given the variety of annotation types in our set (*e.g.,* aspects, species), we used a measure that counts the number of descendants (*dcnt*), or child terms, for each entry. Higher level terms will have a larger score and are considered general or global. More descriptive terms with no descendants, or leaf-level terms, are more precise or detailed and receive the lowest *dcnt* value. The *dcnt* analysis quantitatively demonstrates that CACAO annotations are made to specific terms (Fig. 2C). That pattern is consistent with the way annotations were reviewed, where only the most specific term that could be chosen based on the details reported in the paper was counted correct. For comparison, we performed the same *dcnt* analysis on all manual GO annotations available through 2019. The distributions of *dcnt* values for GO annotations are broader, and statistically different from CACAO within each aspect (Fig. 2C). These data demonstrate that community users can contribute high-quality, precise, and scientifically relevant annotations to GO.

### CACAO community curators enrich ontology development

Over time, GO terms and relationships adapt to reflect research progress(35). Small and large-scale rearrangements result from changes in relationships between GO terms to improve the representation of biological knowledge. Regular updates to the ontology are critical for the database to remain relevant and current. The GO Consortium tracks requests to change the ontology as issues via their GitHub repository accessible on the Helpdesk (http://help.geneontology.org/). CACAO users have submitted >50 tickets via this system, resulting in the creation of 49 new GO terms, many of which now have child terms added by others. Given the diverse literature areas read by community curators, many of these new terms are breaking ground in the ontology. At time of writing, the new terms added based on CACAO feedback had been used >650 times by curators. In addition, at least 14 non-term changes, such as clarified definitions and relationships for current terms, have also occurred. A beneficial, unintended consequence of CACAO is that curators are compelled to resolve issues within the ontology and incorporate new knowledge from areas that are not traditionally covered by model organism databases.

## Discussion

### Community member annotations through CACAO add long-term value to GO

The resources available through the GO ecosystem are among the computational tools most cited by biologists(6). Automatically inferred annotations, those made without curator intervention, are temporary but make up a significant dynamic proportion of the total GO annotations at any given time. However, there are an astonishing >6 million manual GO annotations in the Aug. 2020 release of GO files. The quality of computationally assigned annotations relies on a solid undergirding of manual annotations performed by a dedicated biocuration community(36). The efforts described here are not meant to rival the volume produced by dedicated biocurators, nor are they suggested to replace that organized effort. Instead, we demonstrate how small contributions from many individual community members over time accumulate into a valuable and unique resource. By virtue of its decoupling from the traditional funding model, community curation supplements professional biocuration, especially in under-funded areas(17).

### Targeted crowd-sourcing with attribution makes CACAO annotation sustainable

Recognizing the need to pull expertise from diverse bench scientists, various other initiatives have been implemented to encourage community participation, including asking non-expert ‘crowds’ to help correct the ontology with lower cost and similar accuracy to experts(37). Another natural by-product of this crowdsourcing influx is the diversification of the biocuration workforce. Such introduction of new expertise and perspectives, as is so often the case with trainees, is analogous to the workplace observation that diverse teams innovate and produce more than homogenous ones(38). While the majority in the ‘crowd’ may be unlikely to participate(39), the CACAO implementation of GONUTS is a sustainable model for community contribution of vetted GO annotations in areas of current interest because it caters to a nonrandom crowd, primarily students in an academic course setting.

In a resource-limited environment, the need to incentivize data curation has been creatively approached with different methods such as the micropublication format(40–42). The PomBase community curation project took form as an online annotation tool called Canto where researchers can curate their own publications. Canto has garnered up to an impressive 50% response rates for co-annotation from authors within their community(43). Yet, motivating researchers to weigh in on ontology structure is a long-standing challenge(20). Recognizing the need to credit individuals for their annotation efforts, UniProt now offers a portal for submitting literature-based curation linked to an ORCiD (https://community.uniprot.org/bbsub/bbsub.html)(44), as does the new Generic Online Annotation Tool built for the plant community (http://goat.phoenixbioinformatics.org/). Importantly, the GONUTS wiki provides a web-based public record of CACAO contributions on the website, allowing individuals to cite their efforts.

### CACAO contributions are valuable because they are unique

As NIH-funded resources for microbes of public health importance are being consolidated into broad bioinformatic resource centers, community investment into annotation through a standard pipeline is warranted(45–48). On the one hand, community curators can spend the time to read and extract information from redundant papers (those with information highly similar to already curated literature and conclusions) thus enhancing eukaryotic model organism annotation depth and increasing confidence in existing annotations. On the other hand, community curators sample from a vast literature space outside the typical biocurator’s expertise, expanding overall organism coverage, including in microbial organisms as demonstrated here(49). Because microbial genomes are typically smaller, groups of students can make a major contribution. A significant instance is adding ~50% of all phage GO annotations available in the GO annotation files. CACAO has also spurred updates to ontology relationships. For example, a large rearrangement of biofilm GO terms occurred after CACAO users initiated discussion about their parentage and definitions.

### Community curation through CACAO meets modern open-source research and education goals

With online education thrust to the forefront in this era of the global COVID-19 pandemic, sustainable and authentic education-driven engagement solutions are critically needed(30, 50, 51). Direct individual contributions, community-driven research, and classroom-focused efforts in any number of formats (*e.g.* CACAO, Adopt-a-genome(52)) have been useful in developing student skills and in serving the scientific community. From an educational perspective, the competition format is an engaging format that models real-world scientific skill development with regards to critical reading, iterative editing of a product, and peer review. We hypothesize that this mini biocurator experience may have similar benefits with regards to recruitment, retention, and graduation observed with undergraduate research (53, 54). The biocuration model is highly applicable to scientists and trainees worldwide and complies with FAIR (Findable, Accessible, Interoperable, Reusable(55)) data principles, making its results accessible to all. GO annotation for SARS-CoV-2 and its infection of human cells was immediately pursued to aid strategic planning of the pandemic response (http://geneontology.org/covid-19.html). We appeal to scientists to participate in biocuration efforts through GONUTS, UniProt, or a model organism database/the Alliance of Genome Resources where users can contribute from the comfort of any computer(56).

## Materials and Methods

CACAO competitions for intercollegiate teams are hosted on GONUTS (https://gonuts.tamu.edu). Raw data for all users and every annotation history are maintained by custom extensions to the MediaWiki software used by GONUTS(32). Additional information about competition rules can be found at https://gowiki.tamu.edu/wiki/index.php/Category:CACAO. The data presented here encompass annotations generated from 2010-2018, with expanded taxon information retrieved using the UniProt application programming interface (API) as well as the ETE (v3.1.1) module and various tools from BioPython (v1.74) (57, 58). Summary statistics for CACAO annotations given in Fig. 1 were mined from our local database storage.

Fully correct annotation data are transferred from GONUTS regularly via the current Gene Association File (GAF) or Gene Product Association Data (GPAD) file format, as outlined in GO requirements, directly to the European Bioinformatics Institute’s Protein2GO for incorporation into the complete GO annotation files. All currently included annotations are accessible on GONUTS or via the search engine QuickGO (https://www.ebi.ac.uk/QuickGO/annotations) by filtering for parameter “assigned by” CACAO, and are also provided as a supplementary dataset in GPAD format (Supp Dataset 1) (59).

The 01-01-2020 non IEA GAF (goa_uniprot_all_noiea.gaf.gz) and ontology file (go.obo) were downloaded from http://release.geneontology.org/ for the *dcnt* analysis. Values for *dcnt* were calculated according to GOATOOLS on all manual annotations not assigned by CACAO(34). The Mann-Whitney test with a two-sided p-value was used to compare GO and CACAO *dcnt* distributions within each aspect using SciPy(60, 61).

For the phage analyses, the GAF was filtered into a subset using the following TaxIDs from the NCBI Taxonomy browser: 12333 (unclassified bacterial viruses), 1714267 (*Gammasphaerolipovirus*), 10656 (*Tectiviridae*), 10472 (*Plasmaviridae*), 10659 (*Corticoviridae*), 10841 (*Microviridae*), 10860 (*Inoviridae*), 28883 (Caudovirales), 11989 (*Leviviridae*), and 10877 (*Cystoviridae*).

Changes to the ontology initiated by CACAO users were tallied by searching through the GO issue tracker at GitHub (https://github.com/geneontology/go-ontology/issues) for user handles: @jimhu-tamu, @suzialeksander, @sandyl27, @jrr-cpt, @ivanerill, and/or the query text “CACAO” for open and closed issues, then manually reviewed for accuracy. The final list of GO terms used to calculate the annotations is included as [supplemental file 2]. Matplotlib (v3.1.1) and Seaborn (v0.9.0) were used to generate pie charts, box plots, and bar graphs (62, 63). Figures were compiled and rendered with the open-source program Inkscape 0.92.2.

## Acknowledgments

Funding was provided by the National Institutes of Health (awards GM088849 and GM089636 to J.C.H.) and the National Science Foundation (awards EF-0949351 and DBI-1565146 to J.C.H.) and support for teaching space was provided by the Department of Biochemistry & Biophysics at Texas A&M University. Annotators from across the globe have participated in CACAO competitions, including teams from University College London, University of North Texas, Miami University (Ohio), Penn State University, Michigan State University, North Dakota State University, Hofstra University, Swarthmore College, Houston Baptist University, Mississippi State University, University of Wisconsin-Madison, University of Wisconsin-Parkside, University of Central Florida, Otterbein University, Centenary College of Louisiana, Harvard University, John Brown University, Minnesota State-Morehead, Suffolk University, University of California-Davis, Stanford University, Doane University, Drexel University, James Madison University, Oakland University, Radford University, University of Cincinnati, University of Maryland, Baltimore County, and Virginia Commonwealth University. The contributions of hundreds of student users are proudly acknowledged. We are thankful to colleagues in the Gene Ontology Consortium for their active support and collaboration on this community annotation project. The authors extend an apology to any contributors not named here; however, their participation was foundational to the work and is deeply appreciated as well. This manuscript is dedicated to our beloved co-author, the late Dr. James “Jim” C. Hu, a committed educator, microbial advocate, and invaluable scientific community member.

